# Agent-based modeling reveals impacts of cell adhesion and matrix remodeling on cancer collective cell migration phenotypes

**DOI:** 10.1101/2024.12.23.630172

**Authors:** Temitope O. Benson, Mohammad Aminul Islam, Kailei Liu, Ashlee N. Ford Versypt

## Abstract

Understanding the phenotypic transitions of cancer cells is crucial for elucidating tumor progression mechanisms, particularly the transition from a non-invasive spheroid phenotype to an invasive network phenotype. We developed an agent-based model (ABM) using Compucell3D, an open-source biological simulation software, to investigate how varying biophysical and biochemical parameters influence emerging properties of cellular communities, including cell growth, division, and migration. Our focus was on cell-cell contact adhesion and matrix remodeling effects on cancer cell migration.

We simplified enzymatic remodeling of the extracellular matrix and the subsequent enhancements to cellular chemotaxis or durotaxis as a combined effect of localized cellular secretion of a chemoattractant. By varying the chemoattractant secretion rate and contact adhesion energy, we simulated their effects on cellular behavior and driving the transition from a spheroid phenotype to a network phenotype. The model serves as a digital twin for 3D cancer cell culture, simulating cancer cell growth, division, and invasion over 1 week, validated against published data. The simulations track the emergent morphological and collective phenotype changes using key metrics such as cell circularity and invasion. Our findings indicate that increased chemoattractant secretion enhances the invasiveness of the collective cells, promoting the transition to a network phenotype. Additionally, changing cell-cell contact energy from a strong cell-cell adhesion to a weak cell-cell adhesion affects the compactness of the spheroids, resulting in lower circularity and increased collective cell invasion. Our work advances the understanding of tumor progression by providing insights into the biophysical mechanisms behind invasive cancer cell phenotypic transitions.

## 1. Introduction

Cancer migration and invasion are crucial processes in the development of metastasis. Metastasis, the spread of cancer from its primary location to distant sites, remains one of the most complex and deadly aspects of cancer progression^1–4^. The intricate process of cancer metastasis begins with a single abnormal cell that transforms through a series of developmental stages to form a tumor mass capable of invading new tissues. Metastasis involves sequential steps by which primary tumor cells disseminate, invade surrounding tissue, survive the bloodstream, penetrate secondary organs, and colonize distant sites^1,2^. The biological processes that underlie these events include cellular motility, degradation and remodeling of the extracellular matrix (ECM), and cell signaling pathways that regulate cell survival, growth, division, and proliferation of cancer cells^5–7^.

The journey of these metastatic cells is marked by diverse strategies, from single-cell migration to collective invasion. These steps are further complicated by various phenotypic transitions such as epithelial-mesenchymal transition (EMT), mesenchymal-epithelial transition (MET), amoeboid-mesenchymal transition, and mesenchymal-amoeboid transition ^2,7–9^. Understanding these dynamic processes has challenged researchers for decades, particularly in capturing the interplay of numerous factors such as cell adhesion strength, chemoattractant secretion, and extracellular components (e.g. collagen and fibronectin) that influence the navigation of cancer cells through diverse tissue microenvironments ^10–13^. During cell migration, cells detach from the primary tumor mass, often transitioning into amoeboid or mesenchymal phenotypes, allowing them to navigate independently through surrounding tissues ^13–17^. Collective migration involves coordinated movement where cancer cells maintain cell-cell contact adhesions and move as a cohesive unit. Collective migration, observed in aggressive cancers, is believed to enable more efficient tissue invasion while providing cells with a protective environment that protects them from immune detection and confers resistance to apoptosis ^1,2,18,19^.

Researchers have developed various computational and mathematical models to gain insight into the complex behavior of cancer invasion and metastasis^4,20–29^. These models have become essential tools for studying cancer invasion and metastasis, offering insights into the complex dynamics of tumor progression at multiple scales^30–34^.

Traditional mathematical models, particularly those based on ordinary differential equations (ODEs), have provided valuable information, but often do not address the full complexity of metastasis^31,35^. Continuum models that use partial differential equations (PDEs) have been used to simulate tumor growth and invasion at the tissue level. These models treat the tumor as a continuous medium and account for variables such as cell density, nutrient concentration, and ECM composition^36–40^. PDE-based models are beneficial for studying the effects of biochemical gradients, such as oxygen or nutrient gradients, on tumor growth and invasion^41^. For example, oxygen gradients can influence the formation of hypoxic regions within tumors, promoting more aggressive invasive behavior and EMT^42^. Together with empirical data, these models can provide a predictive framework for understanding how microenvironmental factors drive cancer progression and metastasis ^43–46^.

Agent-based models (ABMs) are widely used to simulate the behavior of individual cancer cells and their interactions with the surrounding microenvironment. These models are beneficial for studying cellular heterogeneity and the effects of varying microenvironmental conditions on tumor invasion^16,47–49^. In an ABM framework, individual cells are treated as agents with defined rules governing their behaviors, such as migration, proliferation, and interaction with other cells and the ECM. By adjusting parameters related to the cellular rules and biochemical and biophysical interactions with the microenvironment, ABMs can replicate key cancer invasion aspects, including individual and collective migration patterns ^29,34,50–53^.

Despite significant progress in mathematical and computational modeling, many open questions remain. Key challenges include understanding how various parameters drive metastasis and identifying critical thresholds for transitions between invasion strategies and phenotypes. Even with advancements in technology, current models often lack sufficient resolution in capturing an important phenotypic transition observed in invasive cancers: the morphological switch from a spherical, cohesive cell mass to a more elongated, network-like structure^4,24,25^. This transition, a hallmark of collective migration, is critical to understanding how cancer cells adapt their morphology to optimize invasion in response to physical and biochemical cues within the microenvironment^54,55^. The absence of this phenotypic transition in most computational models limits the ability to capture the full spectrum of cellular behaviors contributing to invasion and metastasis. Previously, we developed a mathematical and computational model to describe and understand cancer migration involving ECM remodeling. In that work, we examined how lysyl oxidase (LOX) facilitates the movement of cells through a tumor microenvironment undergoing remodeling and cross-linking ^28^. We also developed a hybrid discrete-continuous computational model using the software CompuCell3D to simulate the migration of metastatic cancer cells through the ECM during chemical and physical remodeling. We treated cancer cells as discrete agents, while ECM components, such as collagen fibers and remodeling enzymes, were modeled by partial differential equations^29^.

Additionally, modeling efforts are limited in their ability to represent the role of singlecell dynamics in driving these phenotypic changes. While models of cell proliferation and division can simulate tumor growth at the population level, they often overlook the heterogeneity within the tumor and the local interactions that drive the emergence of invasive phenotypes^56,57^. The transition from single-cell behavior to collective dynamics is critical to understanding the progression from benign tumors to aggressive cancers that exhibit enhanced invasion and metastatic potential. Addressing this gap in modeling can offer new perspectives on how individual cell behaviors contribute to tumor evolution and adaptation during metastasis ^58,59^. In this paper, we develop a computational multiscale model to account for the phenotypic switch from spheroid to an elongated collective cancer cell to understand the heterogeneous behavior of cancer cell lines, e.g., the MDA-MB-231 breast cancer cell line^4^. This is important because of our knowledge base so far, of all the *in silico* and mathematical models of cell behavior, there has not been a computational model that captures both the invasive and non-invasive cell structure phenotype. Most *in silico* models consider individual cell, collective cell migration, epithelial, mesenchymal, MET, and EMT transition migration as a singular question of investigation. Experimental models have looked at spheroids and elongation structure due to TGF-*β* ^60^ and how collagen concentration influences cancer cell migration rather than phenotypic structure. We have not yet seen a computational in silico model that captures such a phenomenon. A PhysiCell model ^61,62^ was developed to simulate tumor growth and spheroids formation from the experimental model in ^60,63^ but did not capture elongated/strand formations. Our model, developed using Compucell3D, captures cell growth, division, and elongation formation^64^.

## 2. Methods

### 2.1. Computational model digital twin design for 3D cell-culture experiment

In this paper, we built a computational digital twin for the experimental 3D cell culture results of invasive and non-invasive collective cell structures found in Chen et al.^4^. We built a hybrid ABM for cancer cell behavior coupled to a continuous PDE for secreted chemoattractant chemicals. Chen et al. examined single cells from a metastatic human breast cancer cell line embedded in a 3D collagen cell culture environment. The cells were allowed to proliferate and form collective structures over the course of seven days. Chen et al. observed two distinct collective cell migration phenotypes that they called the invasive network and non-invasive spheroid collective cell structures based on the respective morphologies. The invasive networks exhibited high invasion and low circularity index, while the non-invasive spheroid cell structure exhibited low invasion and high circularity. By two and half days, the invasive network structures invaded three times farther than the spheroid structures. Through immunofluorescence staining, Chen et al. observed laminin-5 (LAM5) and type IV collagen (COL4A1) as matrix components secreted by the MDA-MB-231 cancer cell line. To simplify our computational analysis, we combine the effects of secreted LAM5 and COL4A1 as chemoattractant secretion concentrations.

We developed a computational model that enables users to define parameters to simulate different collective cell structures from a single cell in the center of the simulation environment. We focused our simulation on spheroid and network morphological structures observed in MDA-MB-231 breast cancer cells by Chen et al.^4^. A schematic representation of the experimental conditions simulated is presented in Figure 1. Although our model is based on the experimental study found in Chen et al. ^4^, we aimed to develop an *in silico* model that can be extended to understand mechanisms and phenomena for comprehending the techniques by which cancer cells form different collective cell invasion phenotypes. The details of the model are in the rest of this section.

**Figure 1:**
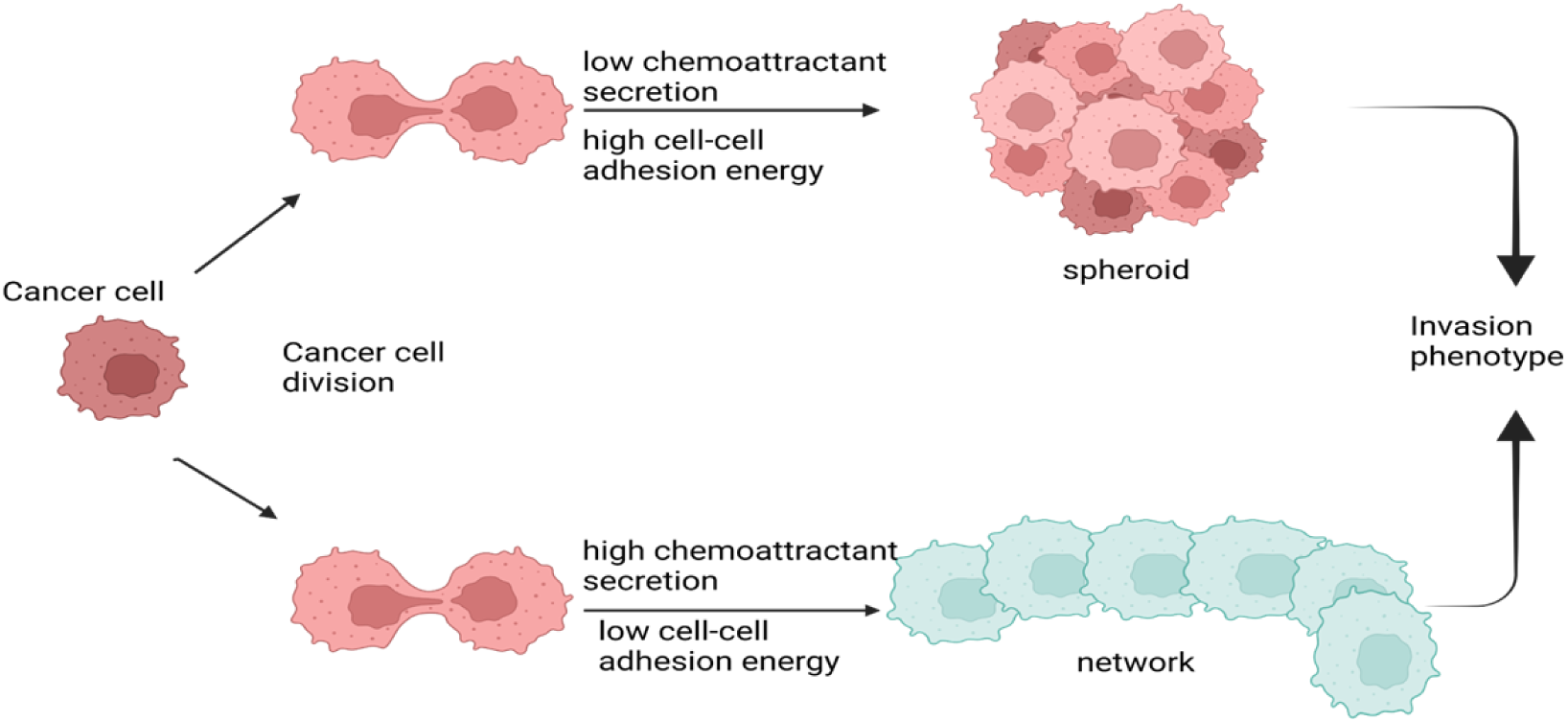
Phenotypic transitions that emerge after cell division of a single MDA-MB-231 breast cancer cell. A single cell within a 3D culture undergoes division, resulting in two daughter cells. Over time, differences in cell-cell contact energy and chemoattractant secretion rate influence the development of distinct collective cell migratory phenotypes—one retaining the compact, cohesive spheroid-like behavior and the other adopting an elongated, invasive network phenotype—highlighting the dynamic plasticity in cancer cell migration. Created with BioRender.com.

### 2.2. Modeling environment

Understanding the fundamental processes that govern cellular behavior is essential for unraveling the mysteries of cellular biology. From the development of organisms to the progression of disease, the behavior of individual cells plays a vital role in shaping the dynamic landscape of living systems. However, studying these phenomena in the laboratory can be challenging due to the complexity and scale of the biological processes. Compucell3D (CC3D) is a multiscale modeling open-source software that combines the lattice-based Cellular Potts model (CPM) with the Glazier-Graner-Hogeweg (GGH) algorithm to simulate the spatiotemporal dynamics within a biological milieu wherein cellular growth, proliferation, invasion, and morphogenesis occur simultaneously^64^. CC3D offers researchers a virtual laboratory for exploring the dynamics of cellular systems. The GGH algorithm provides an intuitive mathematical formalism to map observed cell behaviors and interactions onto a relatively small set of model parameters, making it attractive both to wet-lab and computational biologists^34,64–66^.

The GGH algorithm is designed to model the collective behavior of cells by calculating an energy function called Hamiltonian at each simulation step. Building on our prior simulation design from ^29^, we developed a model tailored to the growth of tumor spheroids in culture with significant changes to incorporate a single cell capable of cell division and proliferation rather than a tumor cell mass. Below, we outline the modifications made to our original CompuCell3D model in^29^ and discuss the implications of these changes. Our previous model simulated cancerous tumor cells as discrete agents within a defined spatial lattice. Each pixel represented a 2 µm length scale, and each biological cell, approximately 20 µm in diameter, was modeled as a computational cell with a width of 10 pixels. The initial tumor mass was a pixelated circle with 100 µm radius, comprising 69 tumor cells with a targeted cell area of 400 µm^2^ as in ^34,67^. The computational domain was a square lattice of 300 pixels per side, ensuring a 200 µm distance between the tumor’s perimeter and the boundary. The most substantial modification is transitioning from an initial cluster of 69 tumor cells in ^29^ to a single cell that undergoes cell division and proliferation. This change allows us to simulate the dynamic process of cultured tumor growth more realistically, starting from a single cell. The cell volume was adjusted to match the new model requirements, with each computational cell width of 8 pixels for validation with experimental spheroid formation and a cell width of 9 pixels for experimental network cell formation. Each computational cell still represents a biological cell with a diameter of approximately 20 µm, but now the model dynamically adjusts cell volume as cells divide and proliferate. Unlike the previous model, which included a static number of cells, the new model incorporates cell growth, division, and proliferation mechanisms. This addition allows the simulation to capture the growth dynamics of a single cell more accurately.

We retained our previous pixel length scale of 2 µm to maintain consistency in spatial resolution. Maintained at 20 µm diameter and 8 or 9 pixels width to ensure compatibility with biological experimental measurements from Chen et al.^4^. The simulation starts with a single cell at the center of the computational domain, which undergoes division and proliferation to form a tumor mass over time. The square lattice remains 300 pixels per side, ensuring the growing tumor mass has ample space to expand and interact with the environment. Starting from a single cell allows the simulation to capture the dynamics of tumor initiation and growth, including cell division patterns, spatial expansion, and tumor cell structure phenotype. The ability to dynamically adjust cell volume during proliferation provides a more accurate representation of cell growth and the spatial arrangement of cells within the tumor. The parameters for CC3D we use here for validation are summarized in Table 1 and for predicting the impact of cell-cell adhesion and chemoattractant secretion rate Table 2.

**Table 1:** Parameter values for validation used in Compucell3D model of cancer cell migration.

| Properties | Values | Units | Comments |
| --- | --- | --- | --- |
| Length per pixel | 2 | $\mu\text{m}$ | Reference<sup>34</sup> |
| Spheroid target cell area ($8 \times 8$ pixels) | $2.56 \times 10^{-4}$ | $\text{mm}^2$ | Reference<sup>4,34</sup> |
| Network target cell area ($9 \times 9$ pixels) | $3.24 \times 10^{-4}$ | $\text{mm}^2$ | Reference<sup>4,34</sup> |
| Lattice width | 600 | $\mu\text{m}$ | |
| Lattice area | 0.36 | $\text{mm}^2$ | |
| Cell migration speed | $1.1 \times 10^{-6}$ | $\text{mm/s}$ | Reference<sup>41</sup> |
| Time per Monte Carlo step | 86.4 | s | Calculated |
| Number of Monte Carlo step in simulation | 7, 000 |  |  |
| Effective cell membrane fluctuation amplitude for Boltzmann acceptance ($T_m = kT$) | 50 | | Reference<sup>68</sup> |
| Pixel copy neighbor order | 2 |  | Reference<sup>64</sup> |
| Adhesion contact neighbor order | 4 |  | Reference<sup>64</sup> |
| Contact energy ($E$) | | | |
| Medium- Medium | 10 | J | Assumed |
| Medium-Spheroids | 10 | J | Assumed |
| Medium-Networks | 10 | J | Assumed |
| Spheroids-Networks | 10 | J | Assumed |
| Spheroids-Spheroids | 10 | J | Assumed |
| Networks-Networks | 15 | J | Assumed |
| Chemotaxis strength toward medium for Spheroid cell | 500 | J | Baseline of chemotaxis strength |
| Chemotaxis strength toward medium for Network cell | 1000 | J | Baseline of chemotaxis strength |
| Chemotaxis secretion rate for spheroid cell | 0.005 | J | Baseline |
| Chemotaxis secretion rate for network cell | 0.05 | J | Baseline of chemotaxis secretion |
| Global decay constant | 0.3 |  | Calculated |
| Global diffusion constant | 1 |  | Calculated |
| Decay coefficient | 0.0 | J | Set |

**Table 2:** Parameter values used for predictive capability in Compucell3D model of cancer cell migration.

| Properties | Values | Units | Source |
| --- | --- | --- | --- |
| Length per pixel | 2 | $\mu\text{m}$ | Kumar et al.<sup>34</sup> |
| Target cell area ($9 \times 9$ pixels) | $3.24 \times 10^{-4}$ | $\text{mm}^2$ | Chen et al.<sup>4</sup> and Kumar et al.<sup>34</sup> |
| Lattice width | 600 | $\mu\text{m}$ | |
| Lattice area | 0.36 | $\text{mm}^2$ | |
| Cell migration speed | $1.1 \times 10^{-6}$ | $\text{mm/s}$ | Zaman et al.<sup>41</sup> |
| Time per Monte Carlo step | 86.4 | s | Calculated |
| Number of Monte Carlo step in simulation | 7, 000 |  |  |
| Effective cell membrane fluctuation amplitude for Boltzmann acceptance ($T_m = kT$) | 50 | | Swat et al.<sup>68</sup> |
| Pixel copy neighbor order | 2 |  | Swat et al.<sup>64</sup> |
| Adhesion contact neighbor order | 4 |  | Swat et al.<sup>64</sup> |
| Contact energy ($E$) | | | |
| Medium-Medium | 10 | J | Assumed |
| Medium-Spheroids | 10 | J | Assumed |
| Medium-Networks | 10 | J | Assumed |
| Spheroids-Networks | 10 | J | Assumed |
| Spheroids-Spheroids | 1–25 | J | Assumed |
| Networks-Networks | 1–25 | J | Assumed |
| Chemotaxis strength toward medium for spheroid cell | 500 | J | Baseline of chemotaxis strength |
| Chemotaxis strength toward medium for network cell | 1000 | J | Baseline of chemotaxis strength |
| Chemotaxis secretion rate for spheroid cell | 0.001–0.1 | J | Varied |
| Chemotaxis secretion rate for network cell | 0.001–0.1 | J | Varied |
| Global decay constant | 0.3 |  | Calculated |
| Global diffusion constant | 1 |  | Calculated |
| Decay coefficient | 0.0 | J | Set |

#### 2.2.1. Cell division and time scale for Monte Carlo steps

CompuCell3D is a widely used tool for simulating the behavior of biological cells using Monte Carlo simulations. One challenge in these simulations is relating the dimensionless number of Monte Carlo steps (MCS) to real-time. Our approach to determining the time scale for cell division specifically focused on the time required for a cell to double its volume. A single cell was placed at the center of a computational simulation grid. Based on our calibration with experimental conditions, the simulation was run for 7000 Monte Carlo steps (MCS), corresponding to 7 days. Cell division rates were recorded and analyzed to understand the proliferation dynamics. Initially, the single cell at the simulation’s center exhibited a high proliferation rate. However, a noticeable slowdown in cell division was observed as the cell divided and the population increased to two or more cells. This behavior aligns with experimental data ^4^, suggesting that the initial rapid proliferation is due to the unrestricted environment of a single cell. In contrast, increased cell density leads to competition for resources and space, thus slowing down the division rate. CompuCell3D does not inherently provide a way to map MCS to real-time units. We need to establish a correspondence between MCS and real-time to connect the simulations of discrete fields to continuous fields involving physical properties such as diffusivity. A common approach is to match the cell growth, division, and invasion observed in experiments with the simulated growth, division, and invasion in pixels per MCS (pixels/MCS). By matching the simulated cell division with experimental data, we established a time scale of 1 day per 1000 MCS for our CompuCell3D simulations, which consistently matched our previous time scale of 1 MCS to 86.4 seconds. Applying this time scale, we determined that cells in our model double their volume every 1000 MCS, equivalent to 24-36 hours in real-time ^46^. This correspondence allows us to accurately simulate cell division and proliferation, enhancing the biological relevance of our tumor growth model.

Haptotaxis is crucial for cell motility; the directional movement involves motile cells migrating up a gradient of chemoattractant sites, which in our model correspond to the concentrations of chemoattractant secreted. To simulate haptotaxis, we utilize the chemotaxis plugin in CompuCell3D. This plugin calculates changes in the system’s effective energy, Δ*H_chem_* during pixel-copy attempts associated with chemotactic motility. The effective energy Δ*H_chem_* is defined in CompuCell3D as

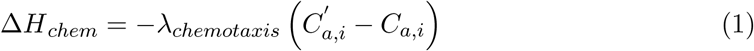

where *λ_chemotaxis_ >* 0 represents the chemotaxis strength, or in our case, the haptotaxis strength *C*^′^*_a,i_, C_a,i_* denote the concentrations of the chemoattractant *C_a_* in the continuous chemical field (determined by a PDE) at the destination and source pixels, respectively, during the pixel-copy attempts. By incorporating the effective energy due to chemotactic motility into the default Hamiltonian of the GGH algorithm in CompuCell3D, the total effective energy of the system, *H*, called the Hamiltonian, calculated for each pixel-copy attempt for pixel *i*, becomes

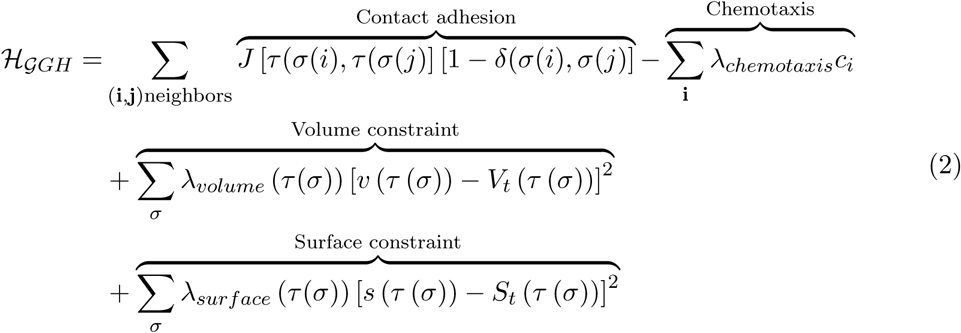

In addition to the chemotactic contribution to the Hamiltonian, we consider contact adhesion, cell surface, and volume constraints.

#### 2.2.2. Contact energies

The contact adhesion energies describe the interaction between neighboring cells, cells, and the ECM. These energies determine how cells adhere to each other and their environment, influencing cell movement, shape, and tissue structure. In CompuCell3D, the contact energy is represented by a matrix, where each entry *J* (*τ* (*σ*(*i*)*, τ* (*σ*(*j*)) indicates the energy between cells *σ*(*i*) and *σ*(*j*). When the contact energy between two cells is high, they are less likely to adhere to each other, leading to less contact and potentially more rounded or separated cells. Low contact energy promotes adhesion, making cells more likely to stick together and form clusters or more cohesive tissues. The overall behavior of a cell can be influenced by the balance of contact energies with its neighbors and the ECM, simulating realistic biological phenomena such as tissue formation and wound healing.

#### 2.2.3. Volume and surface constraints

The volume constraints ensure that cells maintain a realistic size throughout the simulation. Each cell has a target volume and a corresponding volume energy term that penalizes deviations from this target. The target volume is the desired volume of the simulated cell.

The quadratic term [*v* (*τ* (*σ*)) − *V_t_* (*τ* (*σ*))]^2^ penalizes deviations from the target volume, ensuring that cells do not grow too large or shrink excessively. The volume constraint helps accurately model cell growth, division, and death. By maintaining a target volume, the model can simulate biological cells’ physical limitations and mechanical properties.

The surface constraints are similar to volume constraints but apply to the cell’s surface area. These constraints ensure that cells do not become too irregular or overly elongated. The quadratic term penalizes deviations from the target surface area, maintaining the cell’s shape integrity. The volume and surface constraints are defined as

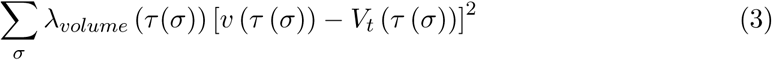

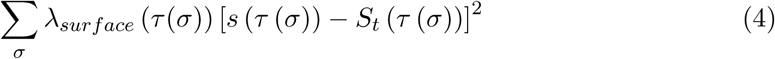

respectively. Maintaining a target surface area is crucial for simulating the physical properties of cells, such as their membrane tension and interaction with the surrounding environment. Equations (3) and (4) describe the surface and volume constraints in the CC3D.

#### 2.2.4. Chemoattractant secretion and reaction-diffusion

The rate of change of the concentration of the chemoattractants *C_a_* per Monte Carlo simulation time steps *t* describes how the conservation of the chemoattractant considering diffusion, first-order decay constant, and secretion processes. This rate of change is given as

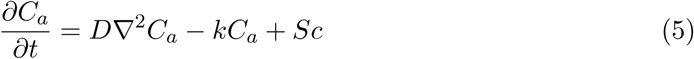

where *c* is the cell source, *D* is the diffusion coefficient, and ∇^2^ is the Laplacian operator that describes the spatial variation of the concentration of the chemoattractant. The combined effect of both *D*∇^2^*C_a_* describes how the chemoattractants spread out over space due to random motion in the simulation environment. The term, −*kC_a_*, accounts for the decay of the substance with a rate constant *k*. The term *S* represents the secretion or production of the chemoattractant substance at a certain rate from the cells.

#### 2.2.5. Cellular growth and proliferation

We developed a mathematical equation using a step function to accurately describe cell growth, division, and proliferation dynamics. Specifically, we generated equations to capture the behavior of cell target volume *V_t_* and the weight of volume constraint *λ_volume_*, which are crucial for understanding how cells adjust their size and replicate over time. We employed the Heaviside step function *H*(*x*) to model these processes. This function allows us to establish conditions for cell growth and proliferation, particularly by specifying the initial target volume for each cell and determining when this volume should double to facilitate division.

The Heaviside step function, *H*(*x*), is defined as

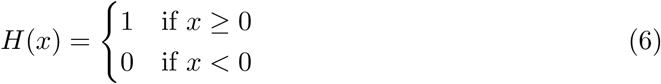

To simulate 3D cell culture experiments and facilitate calibration, we modeled cancer cells to reflect nutrient availability and resource utilization. In this model, a single cancer cell accumulates sufficient nutrients to double its volume by the 700th Monte Carlo (MCS). Subsequently, once the initial doubling occurs, each cancer cell reaches its doubling volume at intervals of 2000 MCS. This approach defines the conditions under which cancer cells grow and divide. The equations governing the cell target volume *V_t_* and lambda volume *λ_volume_* are established as follows, with initial cell volume, *V*_0_ = 51, initial target cell volume doubling time, *t*_0_ = 700 (for a single cell), and subsequent doubling time *t_a_* = 2000 (for multiple cells). These parameters provide the framework for understanding cancer cell growth and division dynamics.

For a single cell (*c* = 1):

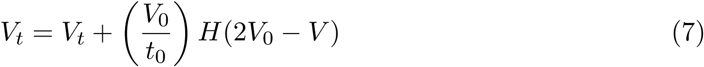

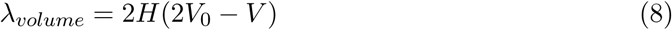

and for multiple cells (*c >* 1):

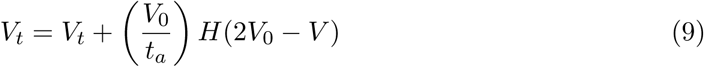

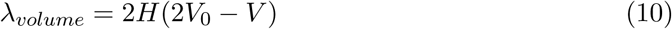

The updated equations, incorporating all conditions above for cell growth, division, and proliferation, are defined as

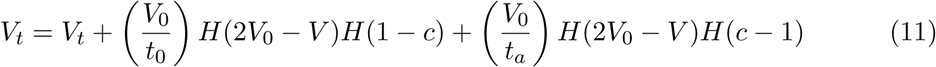

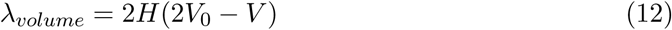

where *V* is the volume of a cancer cell, *V_t_* is the target volume of the cell, *λ_volume_* is the lambda volume of a cell, and *c* is the cell count. Equations (6)–(12) precisely represent the target volume adjustments and the doubling mechanism necessary for cell division. Using the Heaviside function enables clear distinctions between the conditions under which a single cell or multiple cells grow, ensuring that the proliferation dynamics of the experimental result^4^ are accurately modeled.

### 2.3. Parameter scan

The parameter scan feature in CC3D was utilized to explore the input variables’ parameter space systematically. While developing our computational model, we extensively experimented with various parameter configurations to identify optimal settings that align with experimental validation. Specifically, we examined the effects of varying contact energy, cell elongation associated with the network phenotype, chemoattractant secretion rates, neighbor order for contact energies, and parameters within the Cellular Potts model environment. This iterative approach facilitated a robust understanding of parameter sensitivities and their impact on model behavior.

### 2.4. Data extraction and calibration

Accurately calibrating images and cell size is essential for designing and executing computational experiments effectively in computational cell biology. Among the array of software available for this purpose, ImageJ^69^ stands out as a versatile and powerful tool that we used to precisely calibrate images and cells to set up our computational experiments. ImageJ is an open-source image-processing software with rich features for analyzing and manipulating images. One of its key functionalities and features lies in its ability to calibrate images accurately and ensure the measurements made on the images correspond to real-world dimensions. This calibration process involves establishing a scale factor based on known reference points in the image.

The first step in calibrating our experimental image results using ImageJ was acquiring images of reference standards with known dimensions, such as calibration slides or stage micrometers. These reference standards were captured under the same imaging conditions as the experimental samples found in ^4^. Once we obtained the reference images, ImageJ allows for measuring distances and areas between known reference points. ImageJ calculates the scale factor required to convert pixel measurements into physical units by inputting the actual dimensions of these reference points. With this, we obtained physical units of value that we used in setting up our computational experiments accurately and confidently.

### 2.5. Quantification of invasion

Cancer research continually seeks precise and reliable metrics to evaluate the progression and aggressiveness of tumors. Among the myriad of approaches, cell circularity and invasion metrics using the Euclidean distance of cell migration have emerged as significant indicators. These metrics provide insight into the morphological and behavioral changes that occur in cancer cell migration and progression, thereby enhancing our understanding of tumor dynamics and potentially guiding therapeutic strategies ^4,61,70^.

The Euclidean distance, a straight-line measurement between two points, is employed to quantify the extent of cell invasion from the original tumor site. This metric captures the migration of cancer cells, which is crucial for understanding metastatic behavior. *In vitro* assays, such as the Transwell migration assay and the wound healing assay, are commonly used to measure the invasive potential of cancer cells. By tracking the Euclidean distance that cells travel over a specified period, researchers can evaluate the invasive capabilities of different cancer cell lines or the impact of various treatments on cell migration. Higher Euclidean distances indicate greater invasive potential and a higher likelihood of invasion and metastasis. This information can be vital for staging cancer, predicting patient prognosis, and tailoring treatment plans. For example, cancers exhibiting high migratory behavior may require more aggressive treatment strategies, including targeted therapies that inhibit cell motility. The cell invasion is defined as

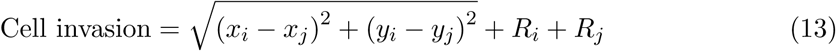

The relevant geometry for this metric is illustrated in Figure 4A.

Cell circularity is a metric used to describe the shape of a cell and allows us to quantify morphological changes. Defined as the ratio of the area of the cell to the area of a circle with the same perimeter, this metric ranges from 0 (indicating an elongated/network phenotype) to 1 (indicating a perfect circle/spheroid phenotype). In the cancer context, cell circularity indicates morphological transformations that often accompany malignant progression. During cancer progression, many epithelial cells undergo EMT, a process where they lose their characteristic shape and become more mesenchymal, exhibiting reduced circularity. This transformation is associated with increased invasiveness and metastatic potential. Therefore, tracking changes in cell circularity can serve as a marker for EMT and the aggressiveness of the cancer. Researchers can quantify cell circularity in tissue samples using imaging techniques and computational software tools. Automated image analysis allows for the high-throughput assessment of cell shapes, providing robust data that can be correlated with clinical outcomes. Lower circularity values typically suggest a more aggressive phenotype, warranting closer clinical attention. The cell circularity is calculated using

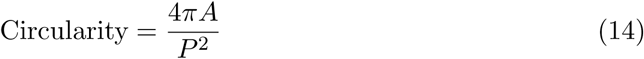

where *A* is the total area of all the cell shapes, and *P* is the perimeter calculated proportionate to the combined diameter of the cell shapes (Figure 2B).

**Figure 2:**
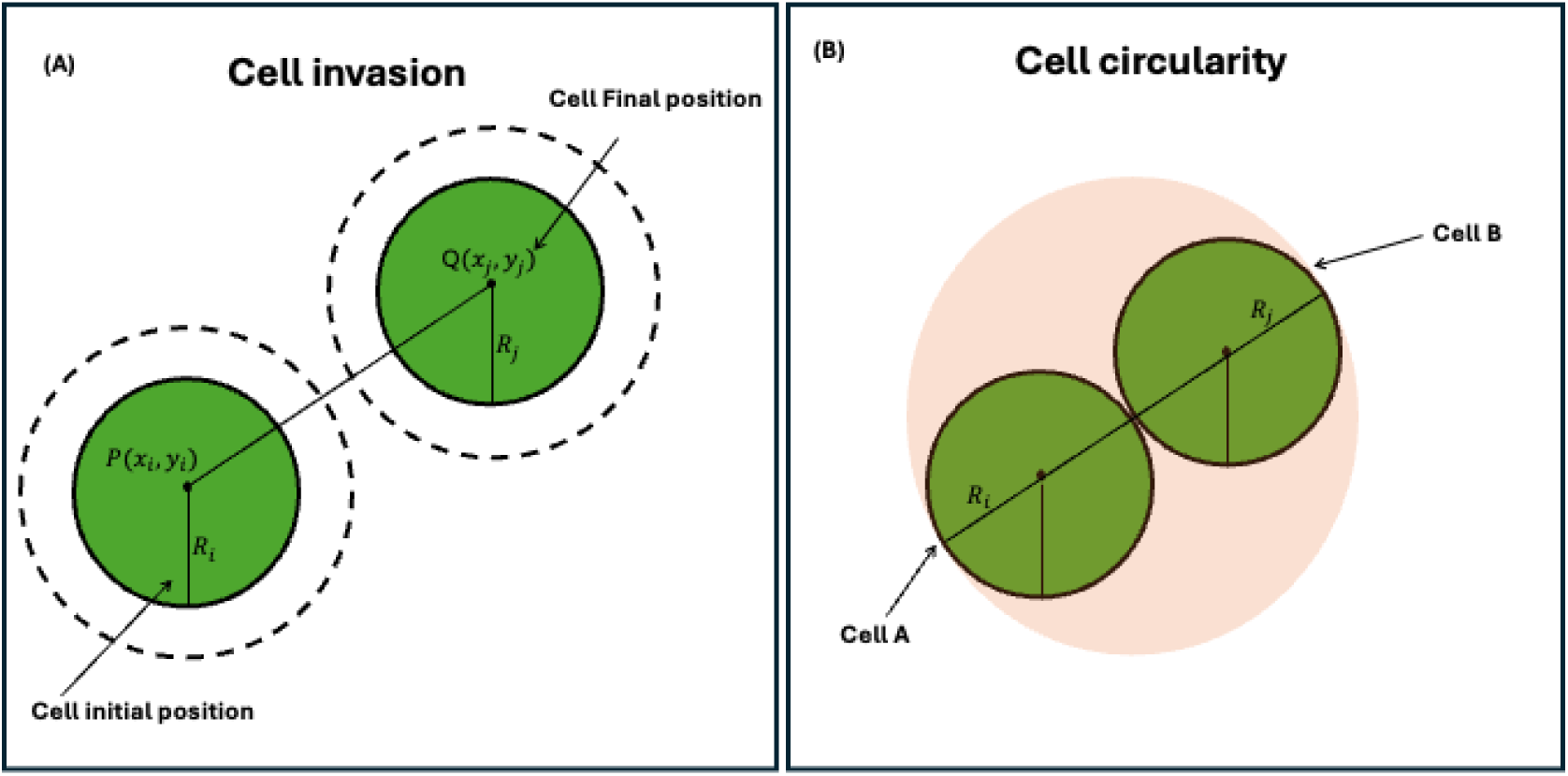
Quantifying cell invasion. (A) Cell invasion is used to measure the distance of each cell from its initial position to the final position. (B) The circularity of two or more cells uses the area of the combined shapes and the perimeter proportion to the longest diameter.

### 2.6. Statistics

All the simulations were replicated 20 times, and data reported in this article are represented as mean ± standard error of the mean (SEM).

## 3. Results

The results begin by analyzing the cell behavior rules assigned to discrete agents within the CompuCell3D framework and validating them with experimental data. We then explore how variations in chemoattractant secretion rates and cell-cell adhesion energies are key determinants shaping invasion phenotypes.

### 3.1. Validation of in silico with experimental results

Our *in silico* models predicted that cancer cells exhibit distinct morphological phenotypes correlating with their invasive potential. Specifically, noninvasive cancer cells were anticipated to form compact spheroids, while invasive cells were expected to develop elongated, network-like structures. To validate these predictions, we used the published experimental studies for cancer cell lines in 3D cell-culture environments^4^. In the experimental assays, the noninvasive cells formed well-defined spheroidal aggregates when embedded in 3D cultures. This is consistent with previous reports highlighting the limited migratory capacity and strong cell-cell adhesion in noninvasive cells, leading to compact structures^71^. In contrast, the invasive cells displayed elongated and interconnected network formations in the same 3D culture conditions. These structures result from enhanced migratory behavior and reduced cell-cell adhesion, facilitating invasion into the surrounding matrix ^53,72^. The cell invasion validation dynamics between the spheroid and network phenotypes are illustrated through three key comparisons. First, we ran 20 simulations for summary statistics for both phenotypes; their mean values are presented at the end of 2.5 days. Second, the cell circularity was compared between simulation and experimental data at the end of day 7. Finally, the maximum cell invasion was analyzed for simulation and experimental results at the end of day 7. These results are summarized in Figure 4. Some sample *in silico* snapshots of the CC3D simulation are provided for both spheroid and network phenotypes at key time points: day 0, pre-division, and days 1, 2, 3, 5, and 7, corresponding to Monte Carlo steps 0, 500, 1000, 2000, 3000, 5000, and 7000 based on our calibration. These CC3D visualization snapshots capture the progression of cell dynamics from day 0 to day 7 and are presented in Figure 3.

**Figure 3:**
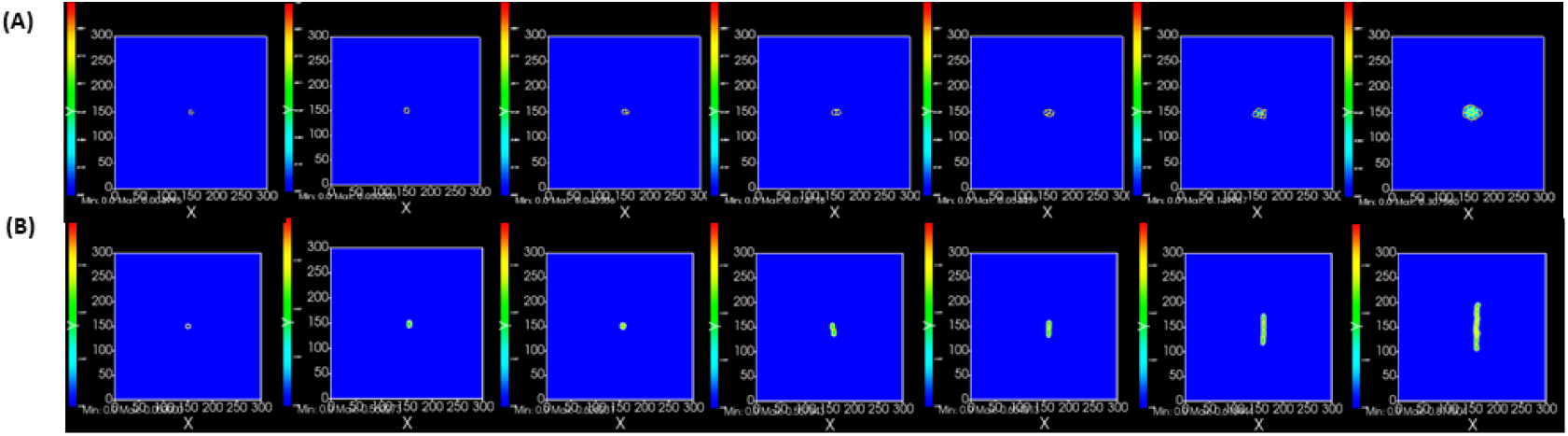
Compucell3D simulation validation dynamics for spheroids and network formation for Monte Carlo time step 0, 500, 1000, 2000, 3000, 5000, and 7000 (left to right).

**Figure 4:**
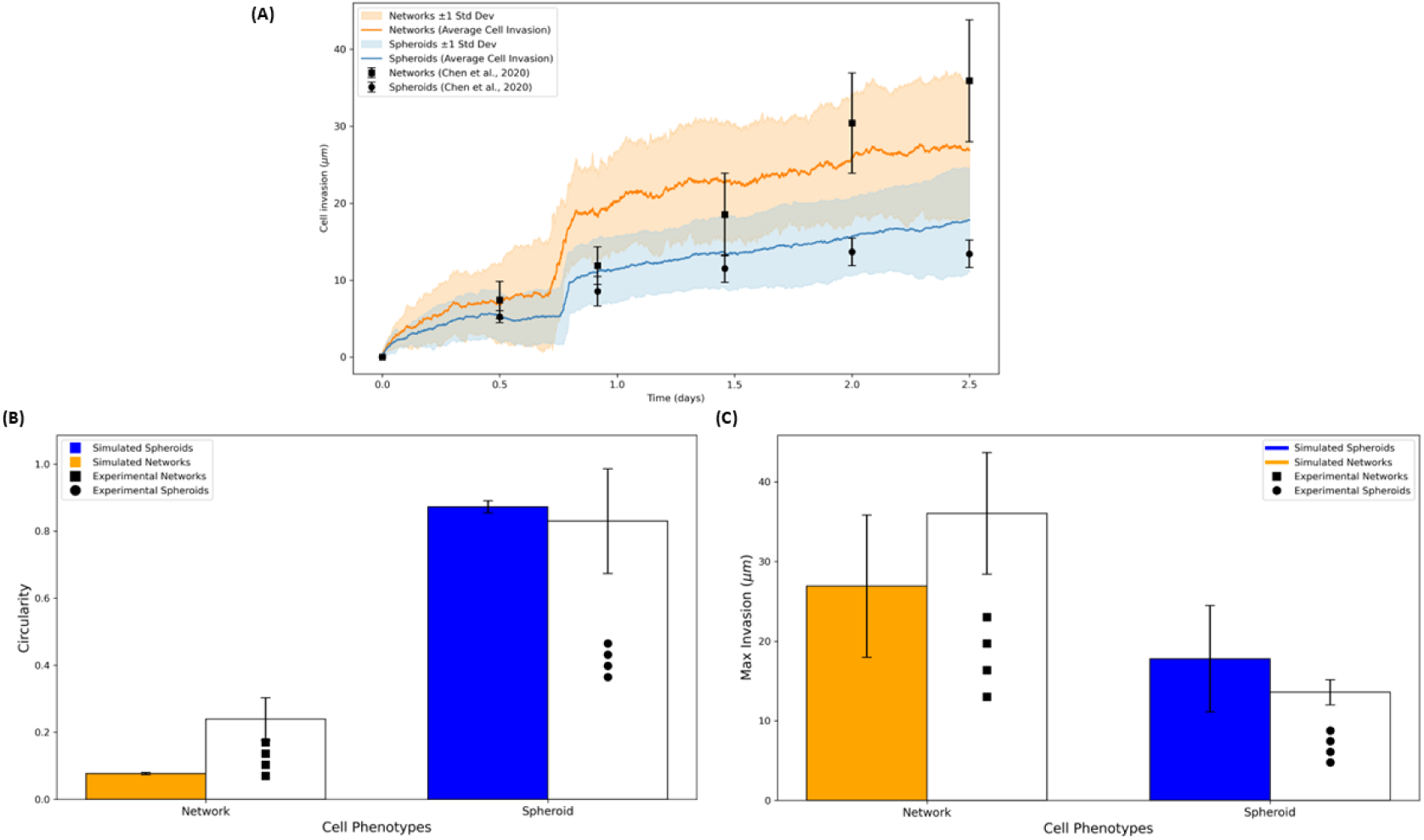
Cell invasion validation dynamics: spheroids and networks phenotype. (A) 20 spheroid and networks simulation results with their mean at the end of 2.5 days. (B) Cell circularity for both the simulation and experiment at the end of day 7. (C) Cell max invasion for both the simulation and experiment at the end of day 7.

Quantitative morphological analysis revealed significant differences between these two phenotypes. Spheroid-forming cells exhibited high circularity and low aspect ratios, indicating their rounded shape. Conversely, network-forming cells showed low circularity and high aspect ratios, reflecting their elongated morphology. These findings align with our *in silico* predictions and suggest a strong correlation between cell morphology and invasive potential. This result can be seen in Figure 4

Collectively, the experimental data corroborate our *in silico* results, confirming that spheroid and network phenotypes in cancer cells indicate noninvasive and invasive behaviors, respectively. This dual approach enhances our understanding of cancer cell morphology’s role in cancer cell invasion and may inform future therapeutic strategies targeting these phenotypic characteristics.

### 3.2. Higher adhesion energy increases circularity and reduces cell invasion in the spheroid cell structure

Our *in silico* experiments also examined the effect of cell-cell adhesion energy on cancer cell morphology, focusing on its role in shaping the invasiveness and circularity of spheroid and network structures. By incrementally increasing cell-cell adhesion energy while keeping other parameters constant, we observed significant changes in cell behavior and structure, reflecting a noninvasive phenotype characterized by high circularity and compact organization. Our result is highlighted in Figure 5.

**Figure 5:**
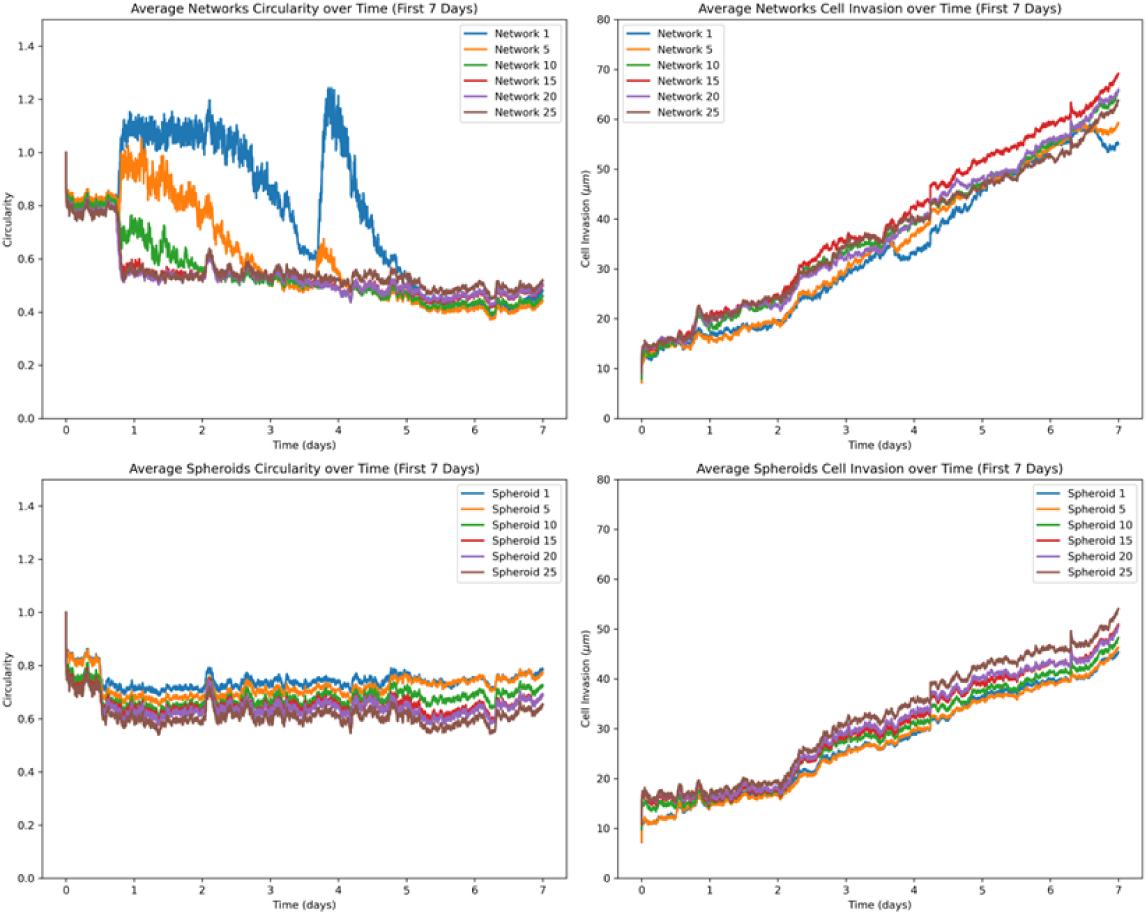
Cell circularity and invasion of *in silico* dynamics for cell-cell adhesion.

At low levels of cell-cell adhesion energy, cells formed loosely organized structures with slight elongation, reflecting a moderate degree of invasiveness. In these conditions, low cell-cell adhesion allowed cells to spread more freely and adopt less compact morphologies, thus reducing overall circularity. This loose arrangement is characteristic of more invasive behavior, where diminished cell cohesion can facilitate migration into surrounding environments ^71^. As the cell-cell adhesion energy increased, the cancer cells displayed higher circularity and became more compact, forming well-defined spheroidal structures. Strong cell-cell interactions promoted tight aggregation at high adhesion energy levels and maintained a round, cohesive structure, limiting individual cell movement and invasion into the surrounding matrix. This morphology indicates reduced invasiveness, as the strong adhesion bonds counteract the cells’ ability to spread and elongate^72^. We vary the cell-cell adhesion energy from a strong adhesion bond to a weaker one in CC3D. In CC3D, a lower value is for a stronger bond, and we vary our adhesion strength in the range of 1–25 and display CC3D snapshots at a particular time in Figure 6 for spheroid phenotype and in Figure 7 for network phenotype.

**Figure 6:**
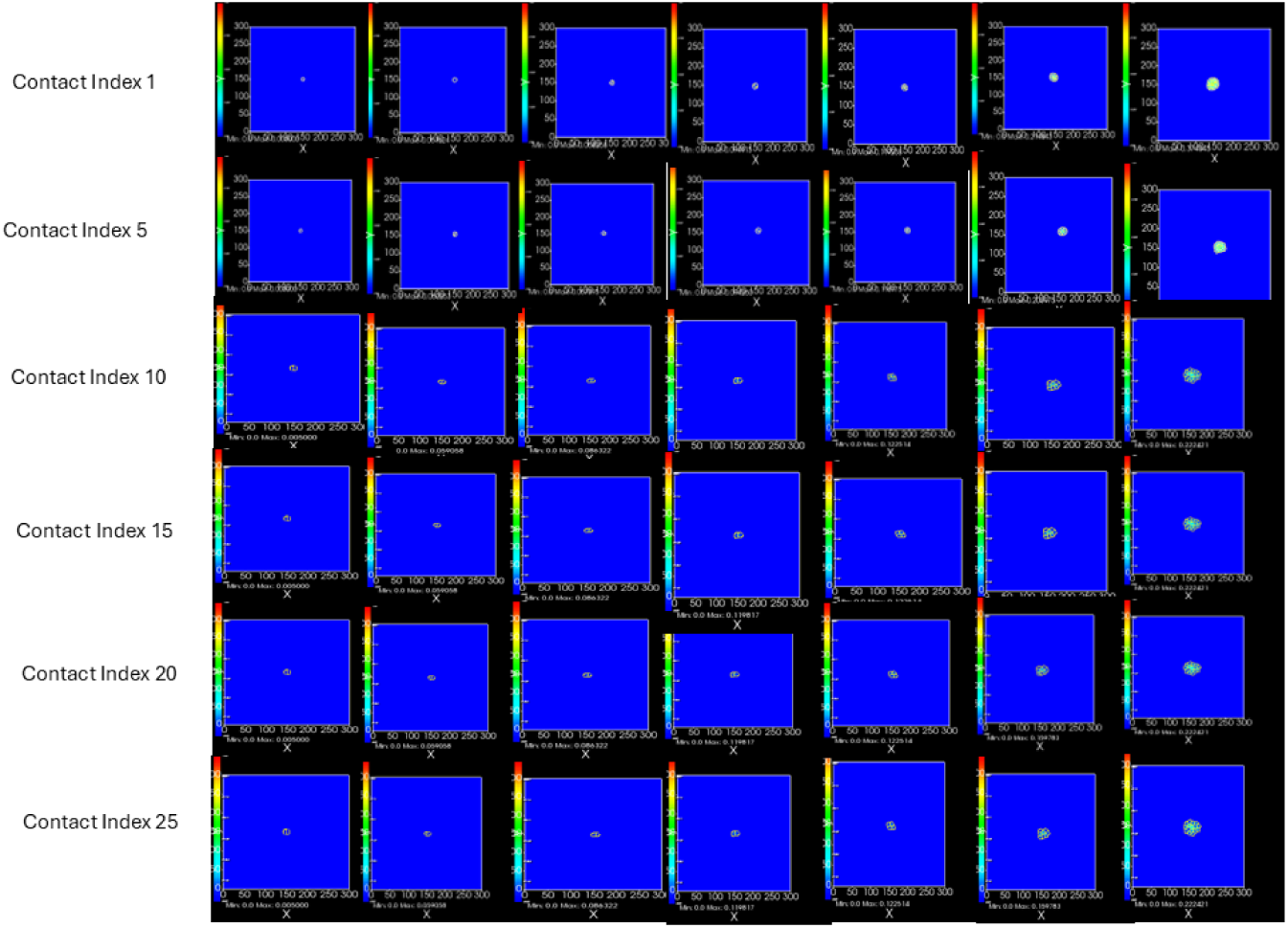
Compucell3D simulation dynamics for spheroids formation for Monte Carlo time steps: 0, 500, 1000, 2000, 3000, 5000, and 7000 (left to right).

**Figure 7:**
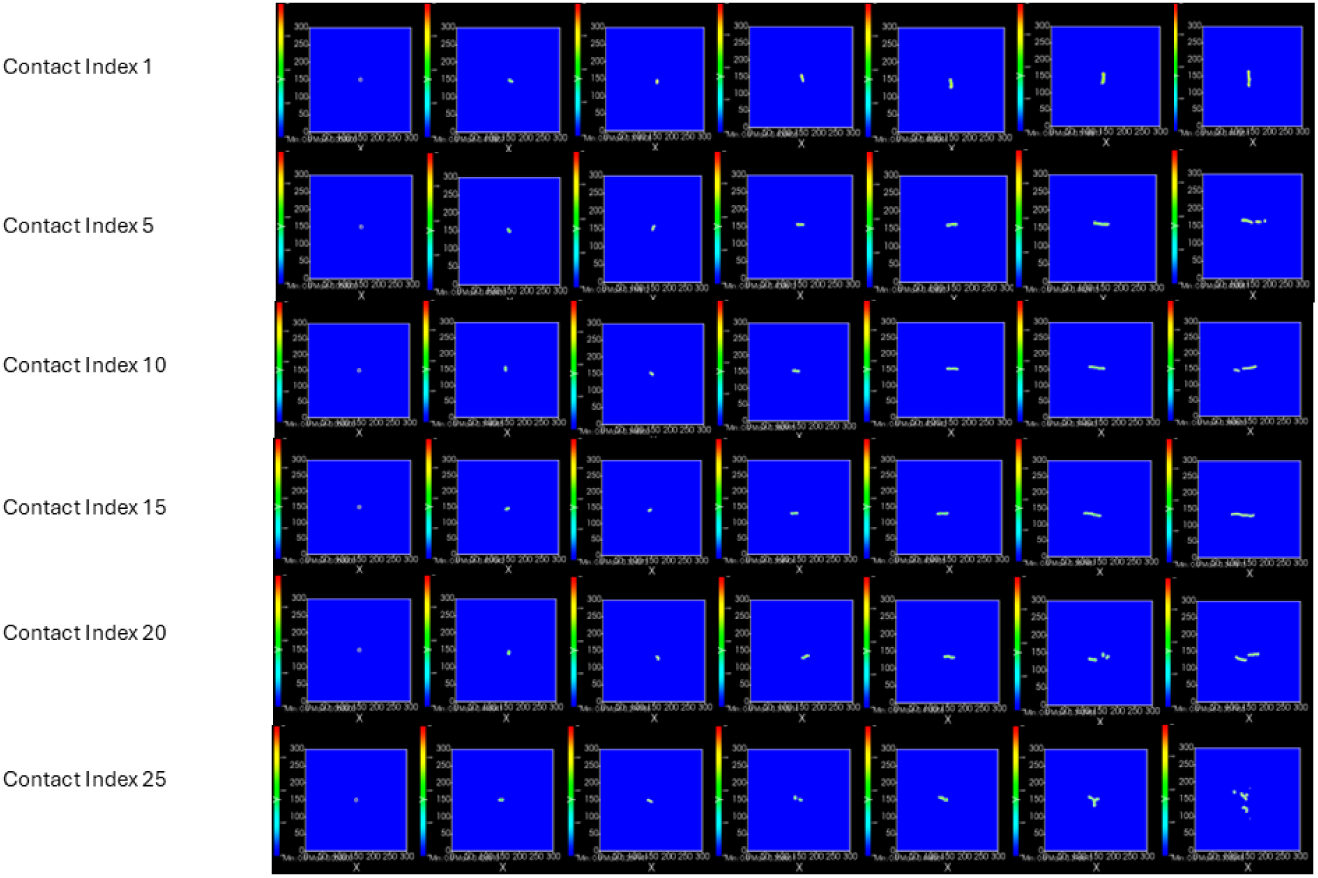
Compucell3D simulation dynamics for network formation for Monte Carlo time steps: 0, 500, 1000, 2000, 3000, 5000, and 7000 (left to right).

Quantitative analysis confirmed a strong correlation between high cell-cell adhesion energy and spheroidal compactness, as evidenced by an increase in circularity and a reduction in the invasion of cell clusters. This shift toward compact spheroids aligns with the characteristics of noninvasive cells, where strong cell-cell adhesion constrains movement and prevents the cells from adopting invasive, elongated shapes. These findings demonstrate that increased cell-cell adhesion energy reinforces spheroidal morphology and is a natural constraint on cell invasion, supporting a noninvasive phenotype^73^.

The simulation results revealed some uncertainty in the circularity of the network cells, particularly under conditions of low cell-cell adhesion energies. This variability is attributed to the inherent stochasticity of the system and the pronounced stretching observed in the network cells. Notably, when these conditions were analyzed manually, the circularity was consistently measured below 1, indicating a more predictable outcome than the simulation. This highlights the model’s weakness to specific parameters and the challenges in capturing deterministic behavior under stochastic influences. Further metrics can be seen in Figure 8, which showcase the impact of cell-cell adhesion on collective cell morphology and invasion.

**Figure 8:**
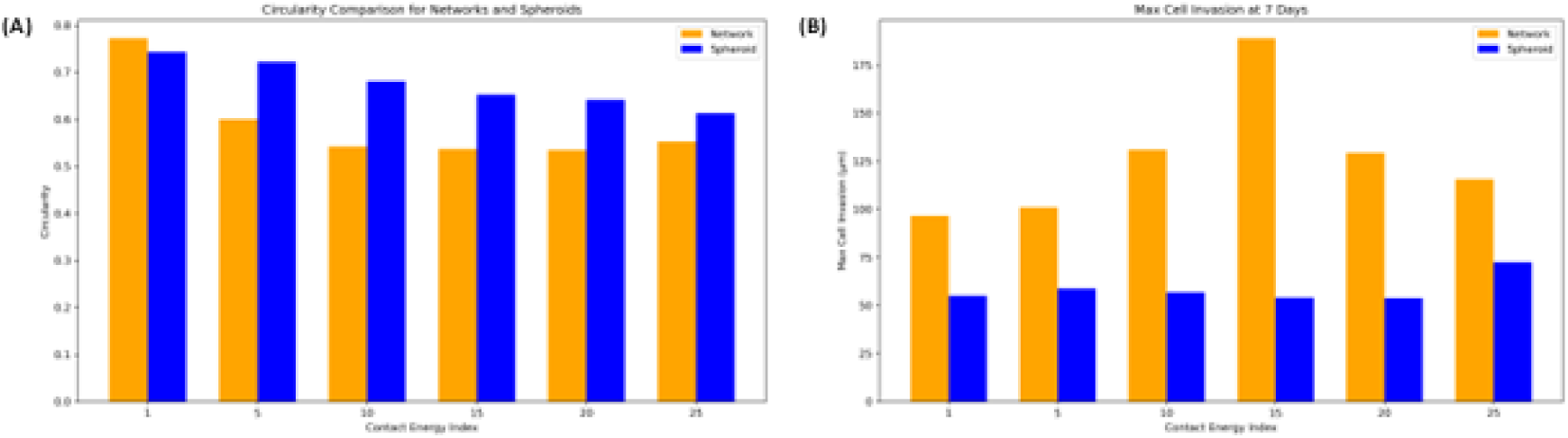
(A) Cell circularity and (B) invasion based on varying cell-cell adhesion strength from strong cell-cell adhesion to weaker adhesion.

### 3.3. Higher chemoattractant secretion rate leads to network structure and increased invasion

We explored the role of chemoattractant secretion rate in influencing cancer cell morphology, specifically its impact on the transition from spheroid to network-like structures. By systematically increasing the chemoattractant secretion rate while holding all other parameters constant, we observed a marked effect on cell shape and organization, indicative of the cells’ invasive potential. At low chemoattractant levels, the cells primarily formed compact, globular clusters consistent with noninvasive behavior. These low-attractant environments were associated with high circularity and minimal elongation, reflecting a cohesive structure where cell-cell adhesion was dominant over motility. This spheroidal morphology aligns with the characteristics of noninvasive cells, as seen in previous studies where limited chemoattractant signals restrict cell migration and maintain a rounded shape ^4,53,71^. Our result is characterized based on cell circularity and invasion in Figure 9.

**Figure 9:**
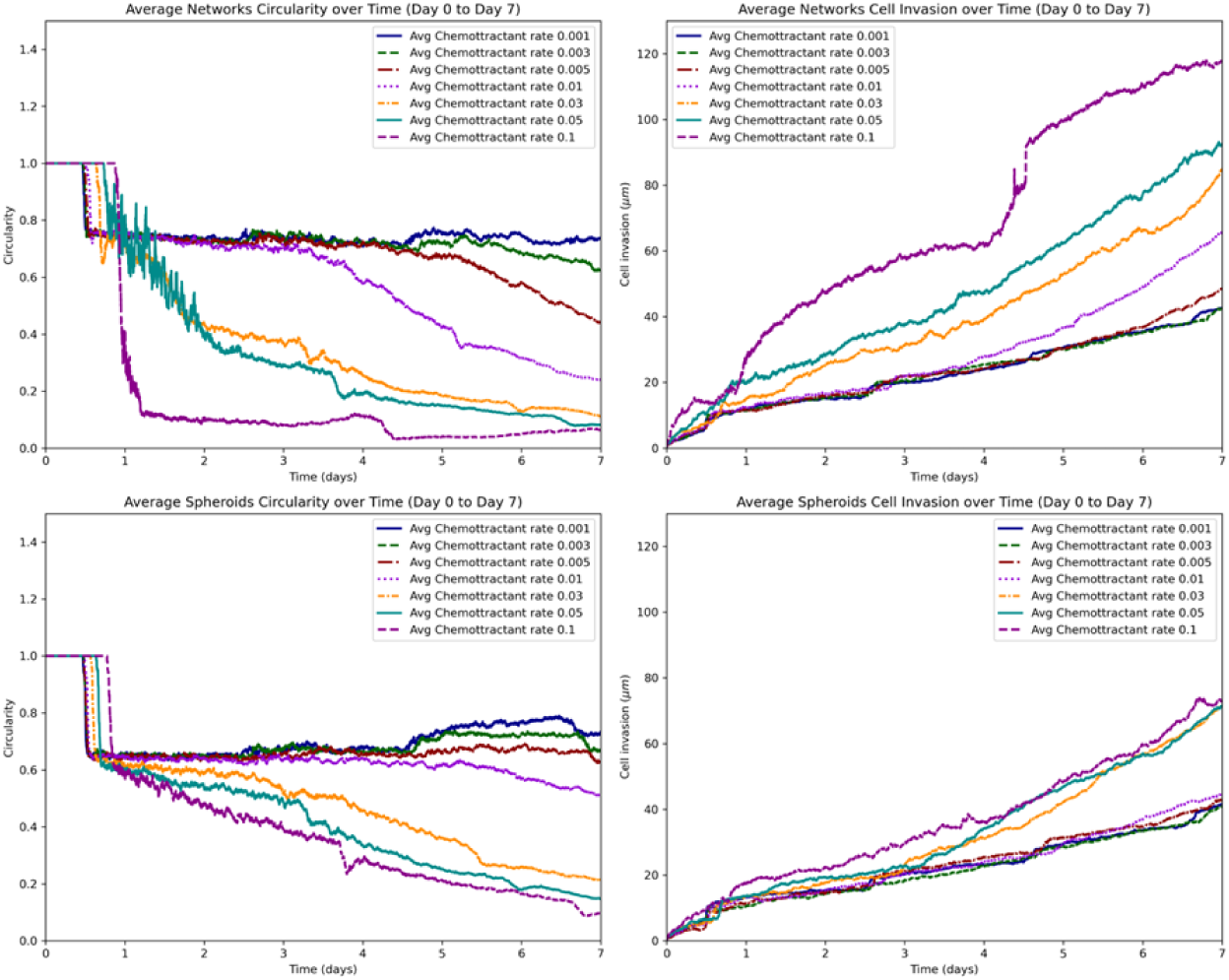
Cell circularity and invasion of *in silico* dynamics for chemoattractant secretion rate.

We observed a distinct morphological transition from round, spheroidal aggregates to elongated, network-like formations in Figure 11 as the chemoattractant secretion rate increased. With moderate increases in chemoattractant, cells began to elongate and form loose clusters. Cells demonstrated significant elongation and interconnectivity at high chemoattractant secretion rates, producing a mesh-like, invasive phenotype characterized by reduced circularity and high cell invasion. This transition from spheroid to network morphology underscores the critical role of chemotactic signals in promoting cell motility and invasion, which enables cells to spread and connect with distant neighbors, facilitating migration and infiltration into surrounding areas ^4,72^. These findings provide evidence that chemoattractant gradients are key drivers of invasive phenotypic behavior, transforming cell morphology from a noninvasive, compact spheroid to an elongated network phenotype as chemoattractant levels rise. This correlation between chemoattractant secretion rate and cell morphology highlights the influence of microenvironmental factors in modulating invasive capabilities in cancer cells, supporting the hypothesis that increased chemoattractant signals enhance invasive phenotypes. Further metrics can be seen in Figure 10, which showcase the transition at the chemoattractant secretion rate of 0.01.

**Figure 10:**
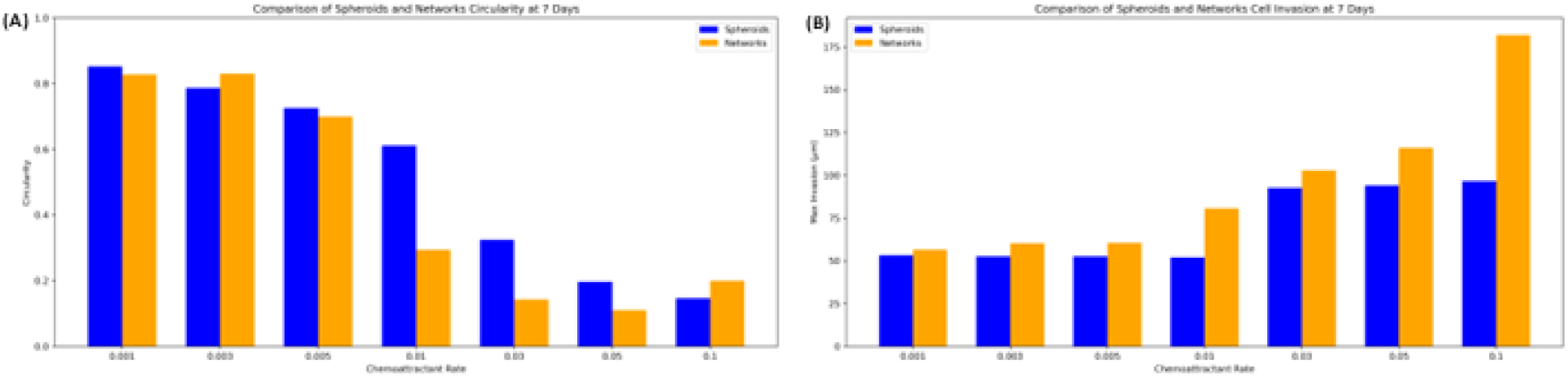
(A) Cell circularity and (B) invasion based on increasing chemoattractant secretion rate.

**Figure 11:**
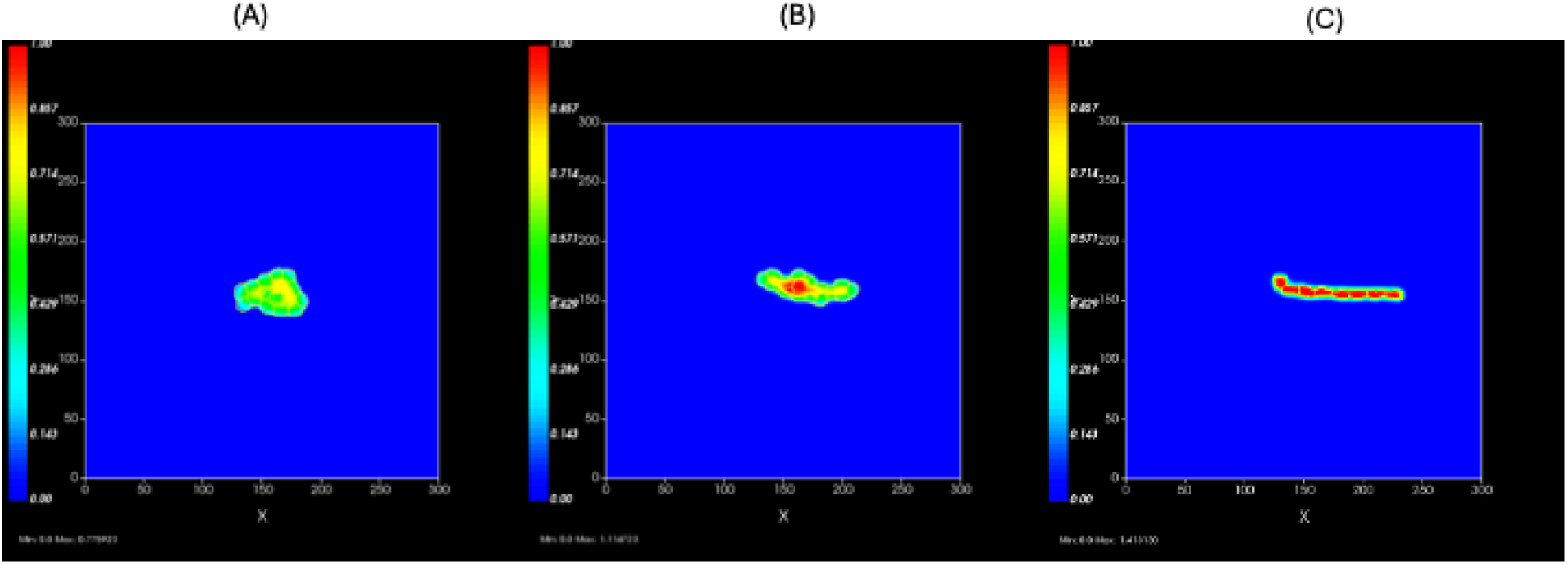
Morphological changes driven by chemoattractant secretion rate from spheroid to network phenotype from a lower secretion value to the higher secretion value (A) 0.001, (B) 0.01, and (C) 0.1.

## 4. Discussion

The results of our *in silico* and experimental validations highlight the critical roles of cell-cell adhesion energy and chemoattractant secretion rate in shaping the morphological and invasive characteristics of cancer cell populations. Our findings show that higher chemoattractant secretion rates correlate strongly with increased cell elongation and network formation, promoting an invasive phenotype. This behavior aligns with existing literature indicating that chemotactic signals drive cellular motility, encouraging cells to migrate and spread into surrounding matrices ^53,72,74^. Conversely, increased cell-cell adhesion energy forms compact, spheroidal structures with high circularity, which limits cell dispersal and reduces invasiveness. This suggests that cell-cell adhesion acts as a physical constraint, encouraging noninvasive, cohesive structures and counteracting the effects of chemotactic forces ^27,62,73^.

These results provide important new information on the biophysical and biochemical processes that underlie the behavior of cancer cells. The interplay between cell-cell adhesion and chemoattractant secretion illustrates the delicate balance of forces that govern cellular morphology and motility. Cells with reduced adhesion but high chemoattractant signals tend to adopt invasive, elongated morphologies, while cells with strong adhesion remain in compact, noninvasive formations. These factors provide a framework for understanding how cancer cells transition between invasive and noninvasive states depending on the molecular and environmental context. With this understanding and enough data, we can build a classification machine learning algorithm that we can combine with the power of an agent-based model for more predictive analysis.

Our findings resonate with prior experimental studies such as those conducted by Gavin et al. ^75^ and Chen et al. ^4^, which investigated brain tumor cells and patient-derived xenograft (PDX) organoids in 3D hydrogels. Gavin et al. observed that ECM composition profoundly influenced organoid growth, viability, and invasion. Using time-lapse microscopy and quantitative image analysis, they demonstrated that Matrigel provided an optimal substratum for local organoid invasion, with behaviors varying across different cell lines and PDXs. Notably, six distinct phenotypic classifications were identified: spheroid and cyst structures were categorized as non-invasive, whereas elongated, protrusive, neuronal, and collective mesenchymal morphologies were deemed invasive phenotypes.

This study’s modifications to our computational parameters have resulted in a more dynamic and accurate CompuCell3D model for simulating tumor cell proliferation, growth, and invasion with improved biological fidelity. By initiating simulations with a single cell and incorporating mechanisms for cell division, our enhanced model more accurately captures the early stages of tumor development and progression. These refinements allow for a biologically relevant representation of tumor dynamics, enabling more profound insights into the mechanisms driving cancer cell morphology, invasion, and growth. This work provides a critical computational tool for investigating tumor behavior and advancing our understanding of cancer progression.

## 5. Conclusions

This study confirms that cancer cell morphology and invasiveness are intricately modulated by cell-cell adhesion energy and chemoattractant secretion. High levels of chemoattractant drive cells toward invasive, network-like phenotypes, whereas increased cell-cell adhesion energy promotes compact, spheroidal structures. These findings highlight the dual roles of chemoattractant-driven motility and adhesion-mediated cohesion in shaping cancer cell behavior and morphology. Through a combination of *in silico* modeling and experimental validation, we have demonstrated that these factors are critical determinants of whether cancer cell populations exhibit invasive or noninvasive properties. This research provides a deeper understanding of the mechanisms underlying cancer cell morphology and identifies potential therapeutic targets to control cancer invasion and metastasis.

Advancing computational and mathematical models to capture the phenotypic transition from spheroid to elongated morphologies will further elucidate the invasion-metastasis cascade. Future models must incorporate parameters such as external potential energies, variations in cell-ECM adhesion, degradation, remodeling, and interactions with the ECM to offer a more comprehensive perspective on cancer invasion and progression. Bridging these gaps will enhance predictive capabilities and inform therapeutic strategies designed to disrupt specific stages of the metastatic process, particularly those targeting the unique vulnerabilities of collectively migrating cancer cell phenotypes. This approach holds promise for improving our understanding of cancer metastasis and developing interventions to mitigate cancer invasion^35,76–78^.

The combination of cell circularity and invasion metrics provides a multifaceted view of cancer progression. While cell circularity offers insights into morphological changes, the Euclidean distance of invasion highlights functional aspects of cell behavior. Integrating these metrics enhances the accuracy of cancer assessments and contributes to personalized medicine. Research has shown that there is often a correlation between decreased circularity and increased invasive distance. Analyzing these metrics together can improve the predictive power of cancer models, facilitating early intervention and better disease management. Advances in imaging technologies and computational analysis are making it increasingly feasible to integrate these metrics into routine clinical practice. High-resolution microscopy and machine learning algorithms can automate the assessment of cell circularity and migration, providing real-time data to inform clinical decisions. Cell circularity and invasion metrics, particularly the Euclidean distance of cell migration, are invaluable tools in cancer studies. By providing detailed insights into the morphological and behavioral changes in cancer cells, these metrics enhance our understanding of tumor dynamics and support the development of targeted therapies. As technological advancements continue to refine these measurements, their integration into clinical practice promises to improve the accuracy of cancer diagnosis and the effectiveness of treatment strategies, ultimately leading to better patient outcomes.

## Acknowledgments

This work was supported by the National Institutes of Health grant R35 GM133763 and the University at Buffalo. We also acknowledge the support provided by the Center for Computational Research at the University at Buffalo: http://hdl.handle.net/10477/79221.

## Notes

### Competing Interest Statement

The authors have declared no competing interest.

